# Modelling erythropoiesis in congenital dyserythropoietic anaemia type I (CDA-I)

**DOI:** 10.1101/744367

**Authors:** Caroline Scott, Damien J. Downes, Jill M. Brown, Christian Babbs, Aude-Anais Olijnik, Matthew Gosden, Robert Beagrie, Ron Schwessinger, Christopher A. Fisher, Anna Rose, David J.P Ferguson, Errin Johnson, Quentin. A Hill, Steven Okoli, Raffaele Renella, Kate Ryan, Marjorie Brand, Jim Hughes, Noemi Roy, Douglas R. Higgs, Veronica J. Buckle

## Abstract

We employ and extensively characterise an *ex vivo* culture system to study terminal erythroid maturation of CD34^+^ progenitors from the peripheral blood of normal individuals and patients with Congenital Dyserythropoietic Anaemia type 1 (CDA-I). Using morphological analysis, FACS analysis and the proteomic approach CyTOF, we analysed patient-derived erythroblasts stage-matched with those from healthy donors during the expansion phase and into early differentiation. In patient cells, aspects of disordered erythropoiesis manifest midway through differentiation, including increased proliferation and changes in the DNA accessibility profile. We also show that cultured erythroblasts from CDA-I patients recapitulate the pathognomic feature of this erythroid disorder with up to 40% of the cells having abnormal ‘spongy’ chromatin morphology by electron microscopy, as well as upregulation of GDF15, a marker of ineffective erythropoiesis. In the tertiary phase of culture, patient cells show significantly less enucleation and there is persistence of earlier erythroid precursors. Furthermore, the enucleation defect appears to be more severe in patients with mutations in *C15orf41*, as compared to the other known causative gene *CDAN1*, indicating a genotype/phenotype correlation in CDA-I. Such erythroblasts are a valuable resource for investigating the pathogenesis of this disease and provide the opportunity for streamlining diagnosis for CDA-I patients and ultimately other forms of unexplained anaemia.

## Introduction

Congenital dyserythropoietic anaemia type 1 (CDA-I) is an autosomal recessive disease associated with characteristic abnormalities in erythroblast nuclei, including binucleate cells and inter-nuclear bridging.^1^ There is ineffective erythropoiesis and macrocytic anaemia, commonly mild to moderate although some patients are transfusion-dependent and the disease can be fatal. A diagnostic feature of CDA-I is the “Swiss-cheese” or spongy pattern of abnormal chromatin in up to 50% of erythroblasts obtained from bone marrow aspirates, visualised using transmission electron microscopy (TEM).^2^ In ~90% of cases, bi-allelic mutations in one of two genes, *CDAN1* (encoding Codanin-1) or *C15orf41* are causative,^3,4^ with the genetic causes of the remaining ~10% of patients yet to be determined. The observation that Codanin-1 and C15orf41 both directly interact with the histone chaperone ASF1^5,6^ implies that the two genes participate in the same pathway. Studies in non-erythroid cells have provided insight into protein function, identifying cell-cycle regulation of *CDAN1*^3,7^ and a potential role in histone supply to the nucleus.^5^ However CDA-I is predominantly an erythroid-restricted disease, and use of erythroblasts for functional analysis would be the most informative way to study CDA-I aetiology and may shed light on how loss-of-function and a range of missense mutations in either gene produces a common erythroid phenotype. Furthermore, as novel *C15orf41* or *CDAN1* variants are identified, pathogenicity cannot be determined without additional diagnostic evidence specific to CDA-I. Availability of cultured erythroblasts for such analysis would avoid the need to perform invasive bone marrow aspirates, particularly important for paediatric patients.

*Ex vivo* systems that closely recapitulate the *in vivo* state provide an important path to understanding the basis of human disease. The expansion and differentiation of CD34^+^ haematopoietic stem cells obtained from peripheral blood has been successfully used to study normal erythropoiesis^8–11^ and to elucidate disease mechanisms in a number of erythroid disorders including Diamond Blackfan Anaemia (DBA)^12,13^ and myelodysplastic syndrome (MDS).^9^ To fully understand the defects that arise in patients who do not generate sufficient mature, lineage-specific cells, any culture system used must recapitulate the terminal stages of erythropoiesis through to enucleation. Furthermore, whilst erythroid culture systems can yield large numbers of erythroblasts for downstream analyses, it has become clear that careful attention must be paid to stage-match diseased and healthy erythroblasts, to ensure findings can be correctly interpreted.^14^

Here we use an *ex vivo* three-phase culture system^8^ (broadly consisting of expansion, differentiation and enucleation) whereby CD34^+^ cells obtained from peripheral blood from both healthy donors and patients with TEM-proven CDA-I are successfully differentiated into reticulocytes. By monitoring differentiating erythroblasts using immunophenotyping by FACS (fluorescence activated cell sorting), morphological assessment, CyTOF (mass cytometry time of flight) and ATAC-seq (Assay for Transposase-Accessible Chromatin using sequencing), we ensure that equivalently staged cell populations are compared. We find patient and healthy erythroblasts comparable during expansion, however, as differentiation proceeds, differences in proliferation rate and alterations in the DNA accessibility profile become apparent. Crucially, at this stage, the pathognomic feature of spongy heterochromatin is recapitulated using this culture system. During the enucleation phase, defects in patient erythroblasts become further evident, with a reduction in enucleation, persistence of precursors and reduced Band 3 expression.

This work shows that appropriate culturing of CD34^+^ erythroid progenitors alongside careful stage-matching of differentiating cells furthers our understanding of CDA-I, and potentially other anaemias that result from ineffective erythropoiesis. Our findings also indicate that such *ex vivo* culturing can be used as a less invasive means to diagnose CDA-I in patients with novel gene variants.

## Methods

Supplemental Tables are available from the authors.

### Study approval, consent and ethics

Subjects were identified through physician-initiated referral through the Oxford Molecular Diagnostic Centre where next generation sequencing was conducted to determine the cause of the patients’ anaemia. Where patients with a molecular diagnosis of CDA-I were identified, patient consent was obtained for entry into this research study. This study was approved by the Wales Research Ethics Committee (REC5) (13/WA/0371) with written consent from patients and/or parents for any samples obtained.

### Isolation of CD34^+^ cells

50ml of whole blood from three healthy donors (two males, one female) and eight CDA-I patients (five with *CDAN1* mutations and three with mutations in *C15orf41*) was collected into EDTA. Full blood counts were performed (Pentra ES60, Horiba) on all samples. Blood was diluted with PBS and overlaid onto Histopaque-1077 (Sigma) and centrifuged for 30 min at 630 *g* (no brake). Harvested peripheral blood mononuclear cells (PBMCs) were washed in PBS and MACS buffer (PBS, 2 mM EDTA, 0.5% BSA) and stained with Human CD34 Microbead kit (Miltenyi Biotec) following the manufacturer’s instruction. The CD34^+^ cell fraction was cryopreserved in freezing media consisting of 90% FBS (Gibco) and 10% DMSO.

### Differentiation of CD34^+^ cells

Differentiation of CD34^+^ cells used a three-phase protocol,^8^ requiring a common base media IMDM (Source BioScience UK) Limited containing 3% (v/v) AB Serum, 10 μg ml^−1^, insulin, 3Uml^−1^ heparin (all from Sigma-Aldrich, Poole, UK), 2% (v/v) fetal bovine serum (Gibco). For differentiation 1×10^5^ frozen CD34^+^ cells were recovered into media containing supplements that are detailed in Figure 1A. Cells were counted throughout.

**Figure 1:**
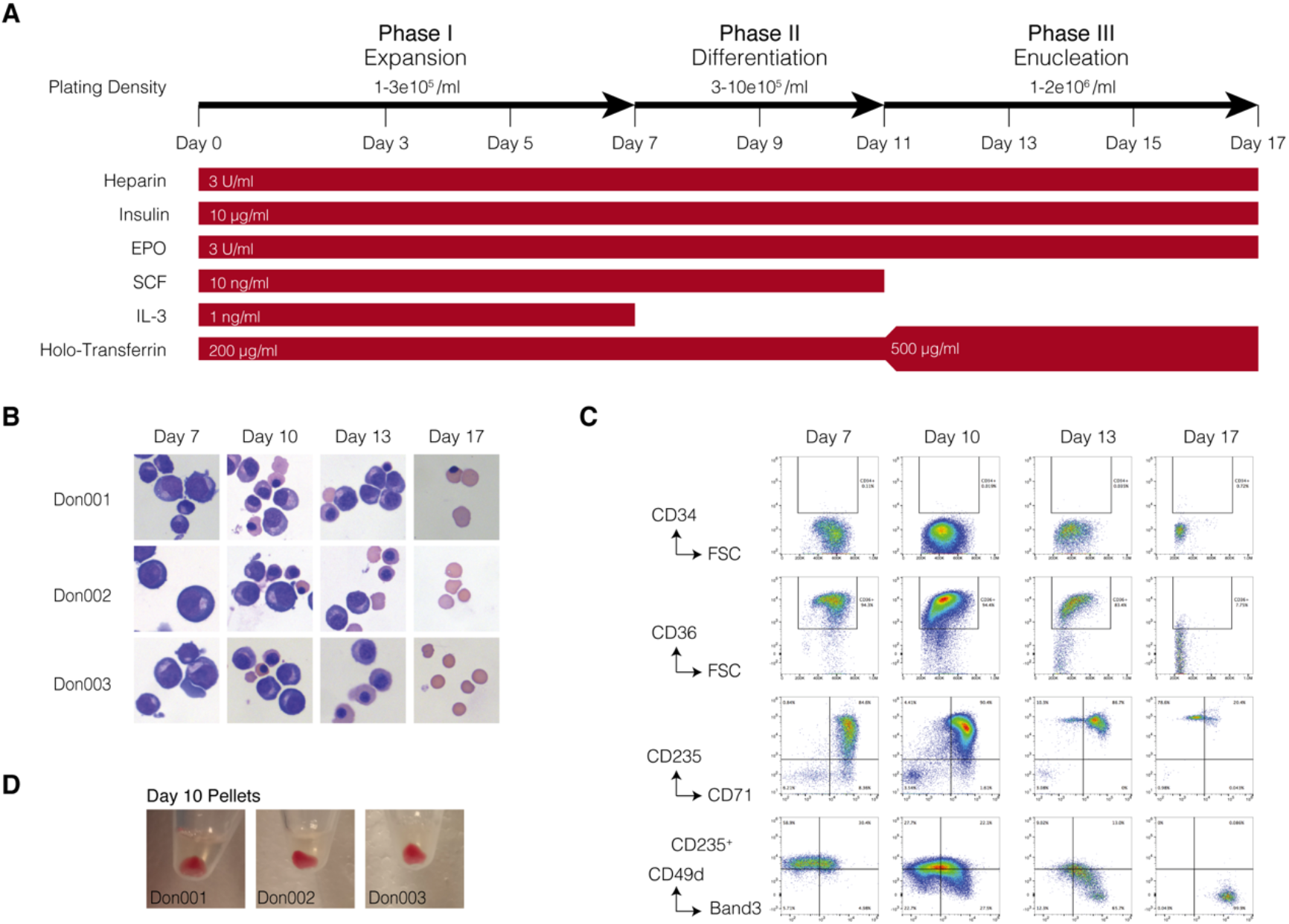
Characterisation of protocol for expansion and differentiation of CD34^+^ progenitors. (A) Schematic of experimental approach for three-phase culture protocol. (B) Representative cytospins stained with modified Wright’s stain (magnification 40x) showing cell morphology during erythroid expansion (day 0-7) differentiation (day 10), enucleation (day 13 & 17). (C) Immunophenotyping of cultured erythroblasts from peripheral blood of healthy donors (n=3) using a 6 colour antibody panel at days 7, 10, 13 & 17 of differentiation. Gates were set using fluorescence minus one (FMO) controls (see supplemental Fig 5). (D) Images of red cell pellets from day 10 cultured erythroblasts.

### Morphological Analysis and Immunophenotyping

1×10^5^ cells were resuspended in 200 μl PBS, spun (5 min, 400 rpm) in a Cytospin 4 (ThermoFisher), stained with modified Wright’s Stain and mounted in DPX (Sigma). Cytospins were imaged using an Olympus BX 60 microscope at 10x and 20x magnification. For FACS analyses 1×10^5^ cells were suspended in FACS buffer (90 % PBS, 10 % FBS) and stained with an erythroid differentiation panel of fluorophore-conjugated antibodies (Supplemental Table 1) and with Hoechst-33258 (ThermoFisher) for live/dead analysis. FACS was carried out on an Attune NxT (ThermoFisher), voltages and compensation were set using Ultra Comp eBeads (ThermoFisher) for antibodies and single stained cells for dyes. Gating was performed using fluorescence-minus-one (FMO) controls (see Supplemental Fig 1). Analysis was performed using FlowJo (v10.4.2). At day 17 enucleation was assessed both by counting the different erythroid subtypes on cytospins and by incubating 1-3×10^5^ cultured erythroblasts with 0.5 μl of 2.5 mg ml^−1^ Hoechst-33342 (ThermoFisher) for 45 minutes at 37°C. FACS then established the number of enucleated cells based on DNA content (Hoechst) and cell size (Supplemental Fig 6A).

### Transmission Electron Microscopy

Approximately 5×10^6^ intermediate erythroblasts staged using morphology and immunophenotyping were prepared for electron microscopy as previously described.^15^ The percentage of nuclei with spongy heterochromatin was determined from large field images.

### Iso-electric Focusing

Approximately 1×10^6^ cultured erythroblasts were lysed in Haemoglobin elution solution and the haemoglobin analysed by iso-electric focusing (IEF) (RESOLVE^®^ Haemoglobin kit, PerkinElmer, USA) using the manufacturer’s instructions on a water-cooled horizontal electrophoresis rig (GE Healthcare). Gels were fixed in 10% trichloroacetic acid and stained with the JB-2 staining system (Perkin Elmer, USA) as per manufacturer instructions.

### RNA isolation and gene expression analysis

1-5×10^6^ cells were fixed in 1 ml TRI-reagent (Sigma), snap frozen on dry ice and stored at −80°C. RNA was extracted by addition of 100 μl 1-bromo-3-chloropropane, separation in a Phase Lock gel Heavy tube (5Prime) and precipitation with 1 μl of GlycoBlue and an equal volume (~500 μl) isopropanol and centrifugation (10 min, 12,000 g, 4°C). RNA pellets were washed with 75% ethanol, resuspended in DEPC-treated water, and stored at −80°C. For RT-qPCR, RNA was treated with 2U of rDNase I (Invitrogen), then 1 μg of RNA was used to generate cDNA using SuperScript III First Strand Synthesis SuperMix (ThermoFisher) following the manufacturers instructions. Real-time RT-qPCR was performed on a StepOne Thermocycler (ThermoFisher) using Taqman Universal PCR Master Mix II (Life Tech) and commercially available assays (Supplemental Table 2).

### Chromatin accessibility

ATAC-seq was performed as previously described^16,17^ using 0.7×10^5^ cells and sequenced on the NextSeq platform (Illumina v2 chemistry) with 39-bp paired-end reads. Reads were mapped to the hg19 genome using NGseqBasic^18^ (V20; --nextera --blacklistFilter --noWindow), which utilises Bowtie.^19^ Sequence depth and mapped reads are provided (Supplemental Table 3). GEO repositories of sorted cell populations (GSE75384) were analysed by the same method. For visualisation PCR-duplicate filtered replicates were merged using Samtools^20^ (v1.3) and converted to bigwig format with minimal smoothing using deepTools^21^ (v2.2.2; bamCoverage --binSize 10 --normalize using RPKM --min Mapping Quality 30). ATAC-seq peaks were called using Macs2^22^ (v2.0/10 callpeak -B -q 0.01). Peak summits from all calls were extended to 550bp and intersected in BEDtools^23^ (v2.25.0), and filtered for high ploidy regions in MIG viewer^24^ and combined and merged (BEDtools merge) to form a collection of all cell types. Non-promoter peaks were defined as being ≥2kb from a transcription start site (TSS). Principal component analysis (PCA) of healthy donor and patient ATAC samples was performed using Deeptools on the top 1,000 peaks. For differentiation trajectory plotting, combined peak calls from sorted hematopoietic populations covering 176,135 non-TSS associated nucleosome depleted regions (Supplemental Tables 4 and 5) were first used to generate an erythroid differentiation trajectory PCA map, onto which samples from differentiation cultures were mapped.

### Immunofluorescence

1-2×10^5^ cells were washed and allowed to settle on poly-L-lysine treated coverslips for 5 min. Cells were fixed in 4% PFA for 15 min and permeabilised in 0.2% Triton X-100 in PBS for 10 min at RT. Slides were blocked using 10% fetal calf serum in PBS at RT for 30 min. Antibodies were prepared in blocking solution at the following concentrations: goat anti-GDF15 (ab-39999; Abcam 1/50) and donkey anti–goat Alexa488 (A-11055; Thermo Fisher Scientific 1/300). Coverslips were mounted in Vectashield with 1 μg ml^−1^ DAPI added as a nuclear counterstain.

### Antibody labeling, barcoding and mass cytometry for CyTOF

Samples were prepared for CyTOF as previously described.^25^ The antibody cocktail used for mass cytometry is shown in Supplemental Table 6.

### Mass cytometry data analyses

Files were processed following Fluidigm recommendations, including randomization and normalization using EQ Beads signal. Files were concatenated, de-barcoded and randomized according to Fluidigm instructions using the CyTOF^®^ Software. FCS files exported from the Helios machine were imported into FlowJo v10. 150,000 events were subsampled from each file (three healthy donors and three *C15orf41* patients). Subsampled events were then concatenated together for Uniform Manifold Approximation and Projection (UMAP).^26^

### Data availability

Sequencing data generated will be available on the Gene Expression Omnibus.

## Results

### Peripheral blood-derived CD34^+^ cells can be differentiated through to erythroblast enucleation

A three-phase *ex vivo* culture system^8^ was used to differentiate erythroid cells from CD34^+^ haematopoietic stem cells (HSCs) derived from peripheral blood of healthy donors (Figure 1A). In addition to the morphological assessment of cultured erythroblasts we characterised their chromatin landscape, globin gene expression profile and the expression of erythroid proteins and transcription factors, to comprehensively evaluate differentiation status. We observed a 1×10^4^ fold expansion of erythroid cell numbers over the course of differentiation. Cellular morphology of normal donor derived cells from pro-erythroblasts to terminal differentiation is shown in Figure 1B. Immunophenotyping revealed a gain of glycophorin A (CD235a) and transferrin receptor (CD71), which typically occurred by day 7 (Figure 1C). As maturation progressed, cells visibly haemoglobinised by day 10 (Figure 1D) and this coincided with increasing expression of the adult globins (Supplementary Figure 2A), with the α-to β-globin ratios remaining around 1 throughout the differentiation (Supplemental Figure 2B). Furthermore, IEF confirmed that during *ex vivo* differentiation predominately adult globin was produced (Supplemental Figure 2C), indicating that erythroblasts undergo definitive erythropoiesis in our culture system. Chromatin accessibility, assessed by ATAC-seq, showed open chromatin at the *HBA1/2*, and *HBB* genes and their associated locus control regions, again indicative of adult erythropoiesis (Supplemental Figure 2D-E). During the third phase of culture, erythroblasts underwent the final stages of differentiation, as measured by increased levels of Band 3 (CD233) and a simultaneous loss of the cytoskeletal protein α-4 integrin (CD49d) within the glycophorin-A positive population (Figure 1C). Enucleated cells were visible on cytospins at this stage (Figure 1B).

### CDA-I patients

To date, ~56 mutations have been reported in *CDAN1* and C15orf41,^1,3,4,27^ although no correlation has been reported between the different mutations within a gene and disease severity. Whilst CDA-I patients are characterised by macrocytic anaemia,^2^ there can be differences in the severity of the disease between individuals, even in those with identical mutations.^28^ We examined eight CDA-I patients in this study (Figure 2A). Patients from our cohort (excluding those receiving regular blood transfusion or venesections) have haemoglobin (Hb) levels and mean cell volumes (MCV) within the normal range (Figure 2B), consistent with ~30% of clinical cases.^2^ These patients do however tend to have higher mean cell haemoglobin (MCH), and a reduced red cell count (RBC) compared to healthy donors (Figure 2B). In one patient (UPID6) with CDA-I, confirmed by TEM, a potentially pathogenic homozygous mutation was identified in *CDAN1* although the allele frequency for this mutation is >1% in specific populations. Data from this patient was included in the *CDAN1* mutation group.

**Figure 2:**
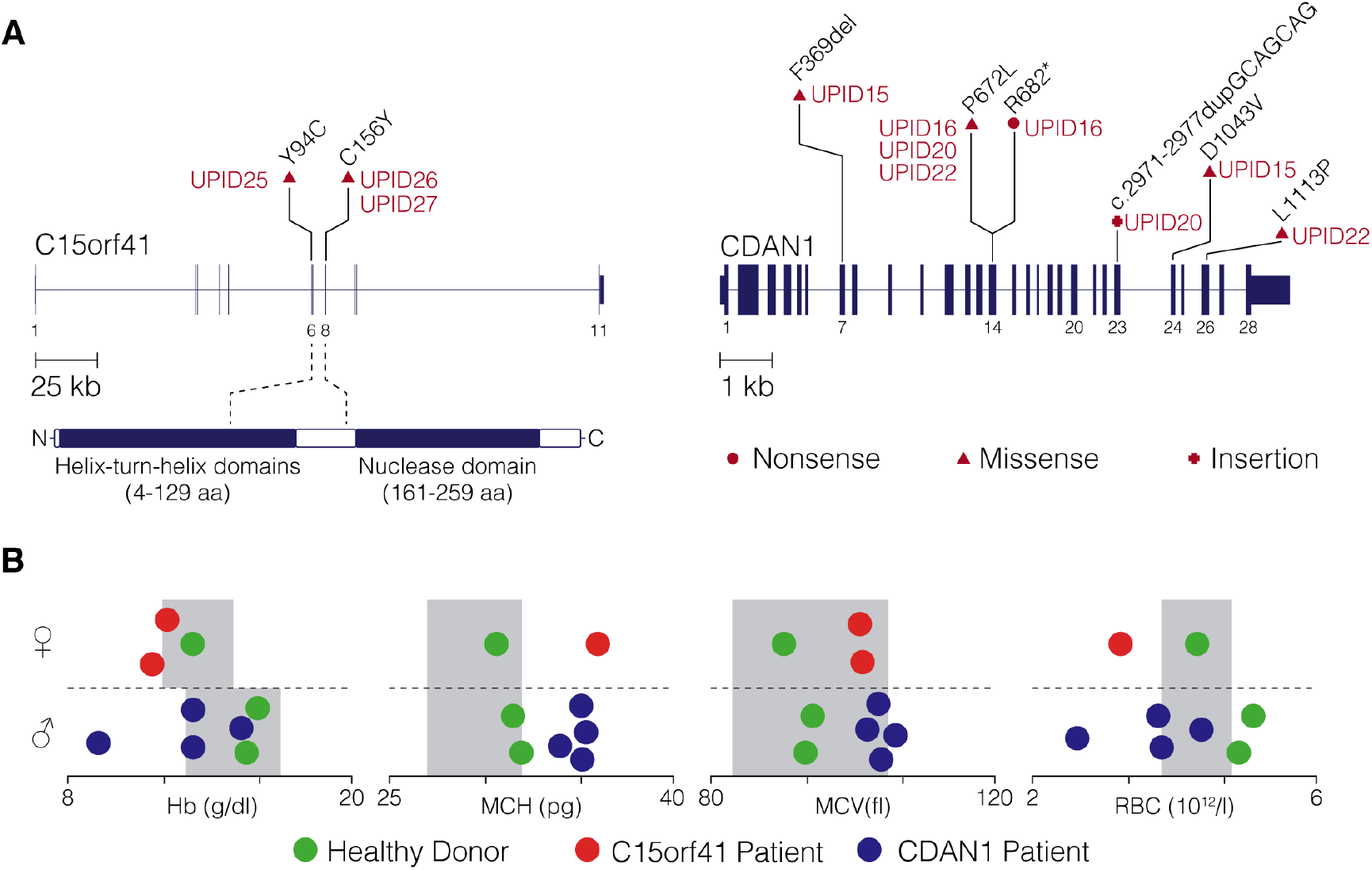
CDA-I patient mutations and clinical data. (A) The location of pathogenic mutations associated with *C15orf41* and *CDAN1* of patients used in this study. Each patient has been given a unique personal identifier (UPID) and the type of mutation indicated. (B) Haematology of healthy donors (green dots) CDA-I patients with *C15orf41* mutations (red dots) and with *CDAN1* mutations (blue dots) where data available. Grey boxes represent the Oxford University Hospital, NHS Foundation Trust Normal ranges for adults. The gender of our cohort is also indicated. Haemoglobin (Hb), mean cell haemoglobin (MCH), mean cell volume (MCV) and red blood cell count (RBC).

### Differentiating erythroblasts from healthy donors and CDA-I patients appear broadly equivalent by immunophenotyping

We studied the expansion and differentiation of erythroblasts derived from CD34^+^ HSCs from CDA-I patients with a variety of mutations in the two known causative genes *C15orf41* and *CDAN1* (Figure 2A). At day 10 the proportions of morphologically identified erythroid cells derived from CDA-I patients with either type of mutation and from healthy donors were similar (Figure 3A and Supplemental Figure 3A). These observations were supported by flow cytometry bulk population analysis showing that differentiation of CDA-I patient HSCs was equivalent to the healthy donors with the simultaneous gain of the erythroid markers CD71 and CD235 (Figure 3B, Supplemental Figure 3A, and for gating strategy Supplemental Figure 1).

**Figure 3:**
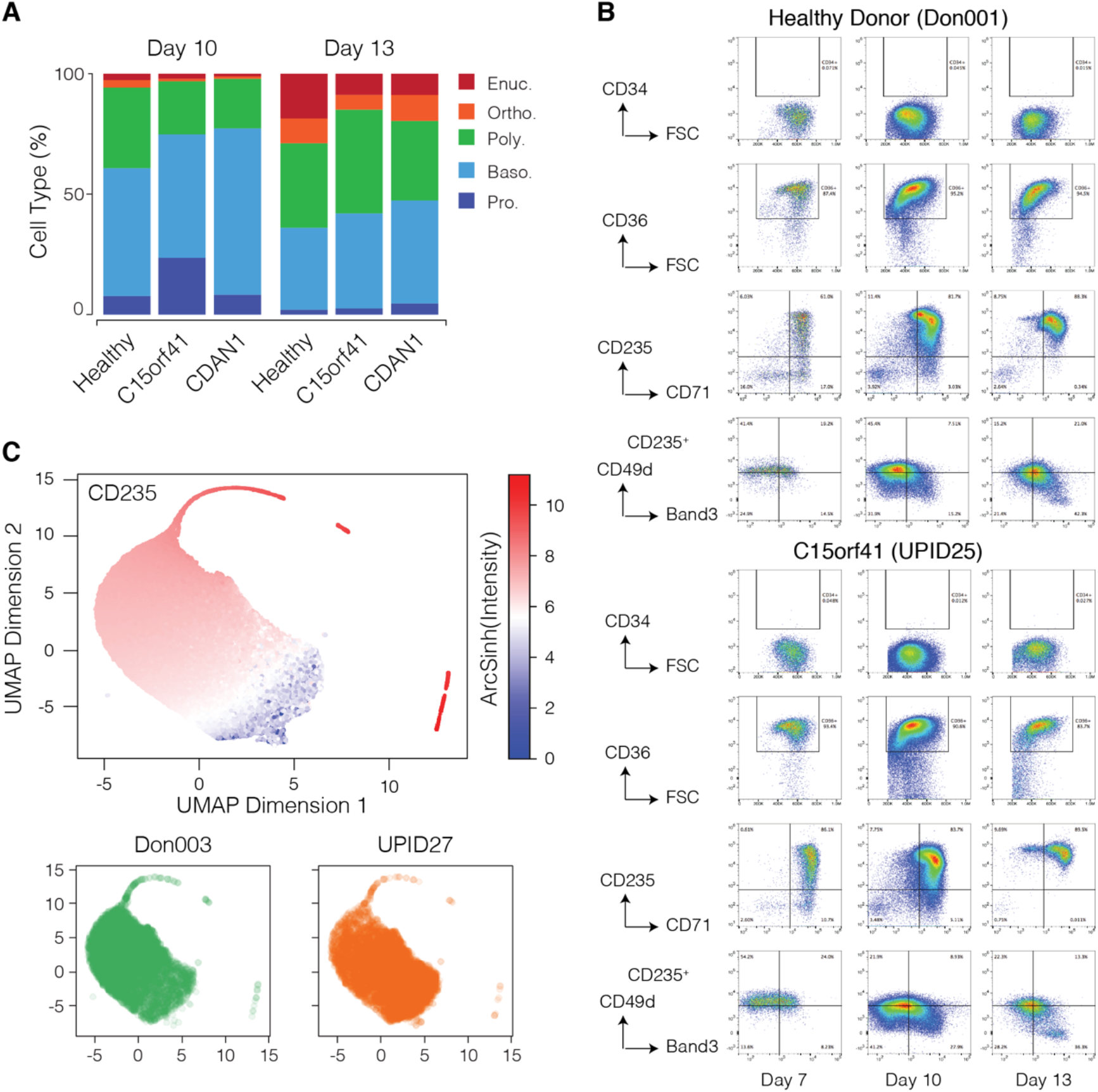
CDA-I patient and normal cultured erythroblasts show similar progression through early erythropoiesis. (A) Percentage of erythroblast stages from cytospins stained with Modified Wright’s stain at two time-points. (B) Representative FACS profiles of cultured erythroblasts from a healthy donor, CDA-I patient with mutations *C15orf41* and CDA-I patient with a mutation in *CDAN1*. (C) UMAP plots showing CYTOF data from healthy donors and *C15orf41*-patient derived erythroblasts at day 11 of differentiation for the erythroid marker CD235 (GPA) (upper panel) and 25 transcription factors and erythroid cell surface markers (lower panel) in a normal donor (Don003) and a CDA-I patient with a mutation in *C15orf41* (UPID 27).

CyTOF is a next-generation flow cytometry platform that allows functional and phenotypic characterisation of cell populations using antibodies conjugated to metal isotopes, thereby avoiding the issues associated with fluorescent overlap encountered using conventional flow cytometry approaches.^29^ Analysis of a panel of erythroid transcription factors and cell surface markers revealed that healthy and *C15orf41* patient-derived erythroblasts follow a continuous trajectory during differentiation (Figure 3C, Supplemental Figure 3B and Supplemental Table 6), with similar levels of CD235 expression at day 11 (Figure 3B, Supplemental Figure 3A). When looking at expression of all erythroid-specific markers analysed by CyTOF there are no clear differences in the clustering pattern between the healthy donors and the patients with *C15orf41* mutations, implying that the erythroid cells from these two sources are phenotypically similar in the early to intermediate stages of erythroid maturation.

### Differentiating erythroblasts from healthy donors and CDA-I patients display defects in chromatin accessibility and organisation

Despite bulk population immunophenotyping indicating the patient and healthy donor cells are grossly stage matched in early differentiation, we observed greater expansion in patient erythroblast numbers compared with healthy donors, which became especially obvious in the later phase of culture (Figure 4A). Such expansion is indicative of ineffective erythropoiesis, which has been characterised as a proliferating pool of immature erythroid cells and sub-optimal production of mature erythroblasts.^30^ Analysis of open chromatin regions can be used to distinguish cell types and deconvolve mixed cell populations;^11,31^ therefore to further compare our differentiating healthy and patient cells we assayed chromatin accessibility using ATAC-seq.^16^ Principal Component Analysis of the top-ranked 1000 open chromatin regions showed a slight differentiation lag in patient cells compared with healthy donors (Supplemental Figure 4). Using a differentiation trajectory of sorted erythropoietic cell populations and 176,135 nucleosome-depleted regions, we mapped both the healthy and patient cell populations against these data. This mapping showed that patient material could be clearly distinguished from healthy donors (Figure 4B), and again suggested an earlier erythroid precursor character to these cells.

**Figure 4:**
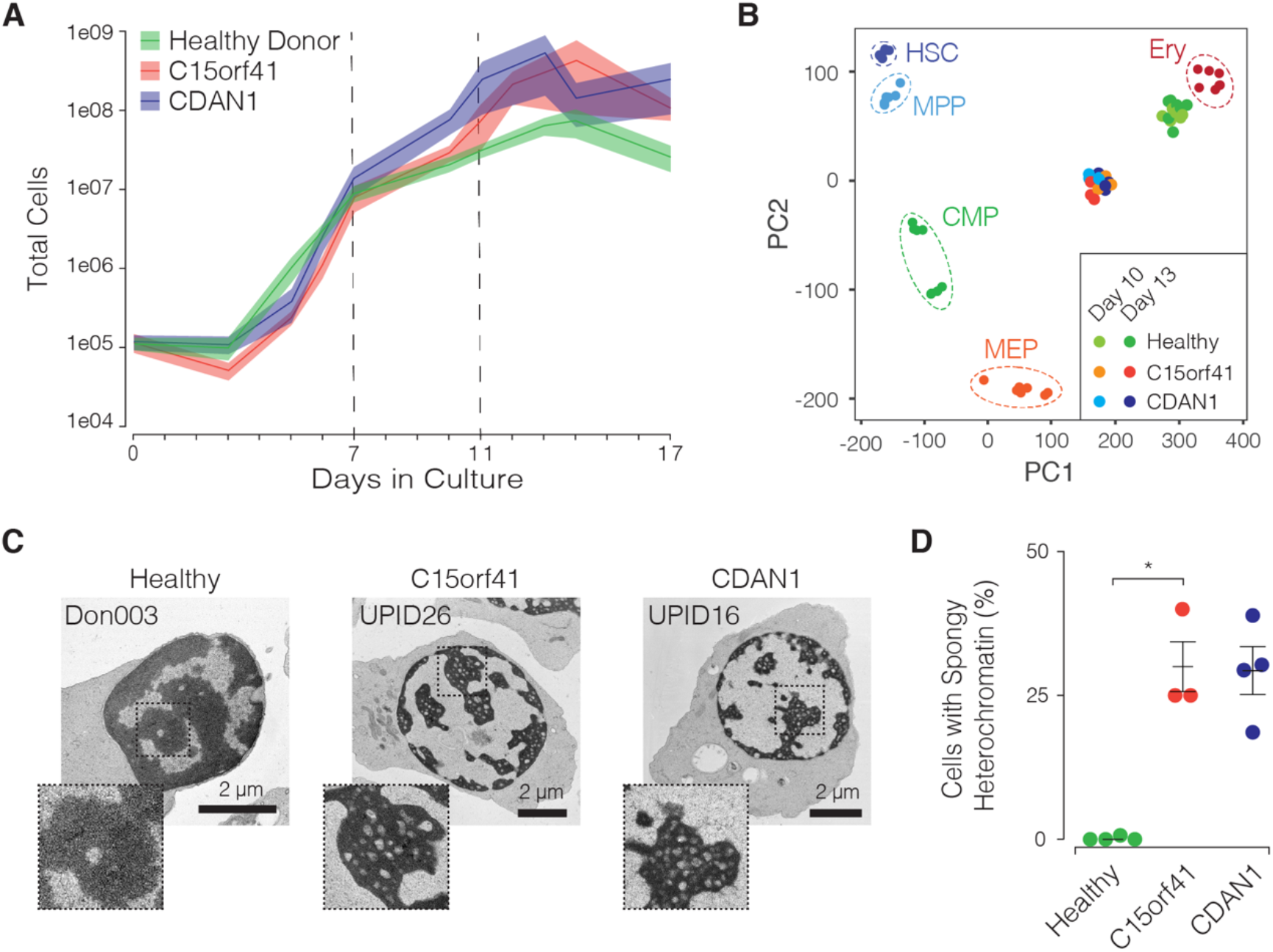
CDA-I cultured erythroblasts show defects in erythropoiesis midway through *ex vivo* differentiation. (A) Proliferation of cultured erythroblasts from healthy donors (n=6), patients with mutations in *C15orf41* (n=3) and patients with mutations in *CDAN1* (n=4) showing increased proliferation in both patient cohorts. Data is shown as mean ±SEM. (B) PCA analysis of ATAC-seq data from *ex vivo* differentiated erythroblasts at two time-points against a trajectory of immunophenotyped progenitors and erythroblasts from bone marrow,^31^ where the patient data is clustered and distinct from healthy donors. (C) Transmission electron microscopy at day 11 of healthy and CDA-I cultured erythroblasts showing the diagnostic phenotype in patients. Inset shows enlarged area to illustrate the pattern of euchromatin and heterochromatin and how this is disrupted in CDA-I patients (D) Percentage of nuclei ±SD with spongy heterochromatin (healthy n= 96/280 and CDA-I n= 96/436 nuclei scored). \**P*=.0189 with Kruskal-Wallis test.

TEM also revealed the pattern of heterochromatin abnormalities characteristic of CDA-I and to date seen only in bone marrow erythroblasts. This feature was observed in all of our patient samples, but not in any healthy donors (Figure 4C and Supplemental Figure 5A), recapitulating the defining disease phenotype. An average of 29% (±7.7 SD) of erythroblasts showed abnormal heterochromatin (Figure 4D). Elevated expression of growth differentiation factor 15 (GDF15), a marker of ineffective erythropoiesis, known to be elevated in serum from CDA-I patients^32^ was also detected by immunofluorescence (IF) in *ex vivo* cultured erythroblasts from CDA-I patients (Supplementary Figure 5B). Therefore, differentiation of CDA-I patient material *ex vivo* successfully recapitulates abnormal patient phenotypes, previously only reported *in vivo*.

### Evidence of a Genotype/Phenotype Correlation in CDA-I

We next assessed the effects of patient mutations on the enucleation stage of differentiation. Firstly, analysis of cellular morphology indicated a persistence of erythroid precursors in CDA-I erythroblasts, particularly in those from patients with bi-allelic *C15orf41* mutations, (Figure 5A), together with a significant reduction in the percentage of enucleated cells (Figure 5B). Levels of enucleation were reduced to a lesser extent in the patients with mutations in *CDAN1* (Figure 5B and Supplemental Figure 6B). Secondly, immunophenotyping of cultured erythroblasts in the enucleation phase revealed changes in Band 3 expression in CDA-I patients (Figure 3B and Supplemental Figure 3A). In normal erythropoiesis Band 3 shows a dramatic increase from the pro-erythroblasts to late erythroblasts,^9^ and while CDA-I patient erythroblasts did progressively gain Band 3, the level of protein was significantly less at day 17 than in healthy donors (Figure 5C). This supports other indications of a delay in the progression of differentiation in patient derived erythroblasts. Notably the Band 3 reduction was more severe in patients with *C15orf41* mutations (n=3 *P*=0.0088) than in the *CDAN1* mutant cells (n=4 *P*=0.0127) (Figure 5D), suggesting the existence of a genotype/phenotype correlation.^33^

**Figure 5:**
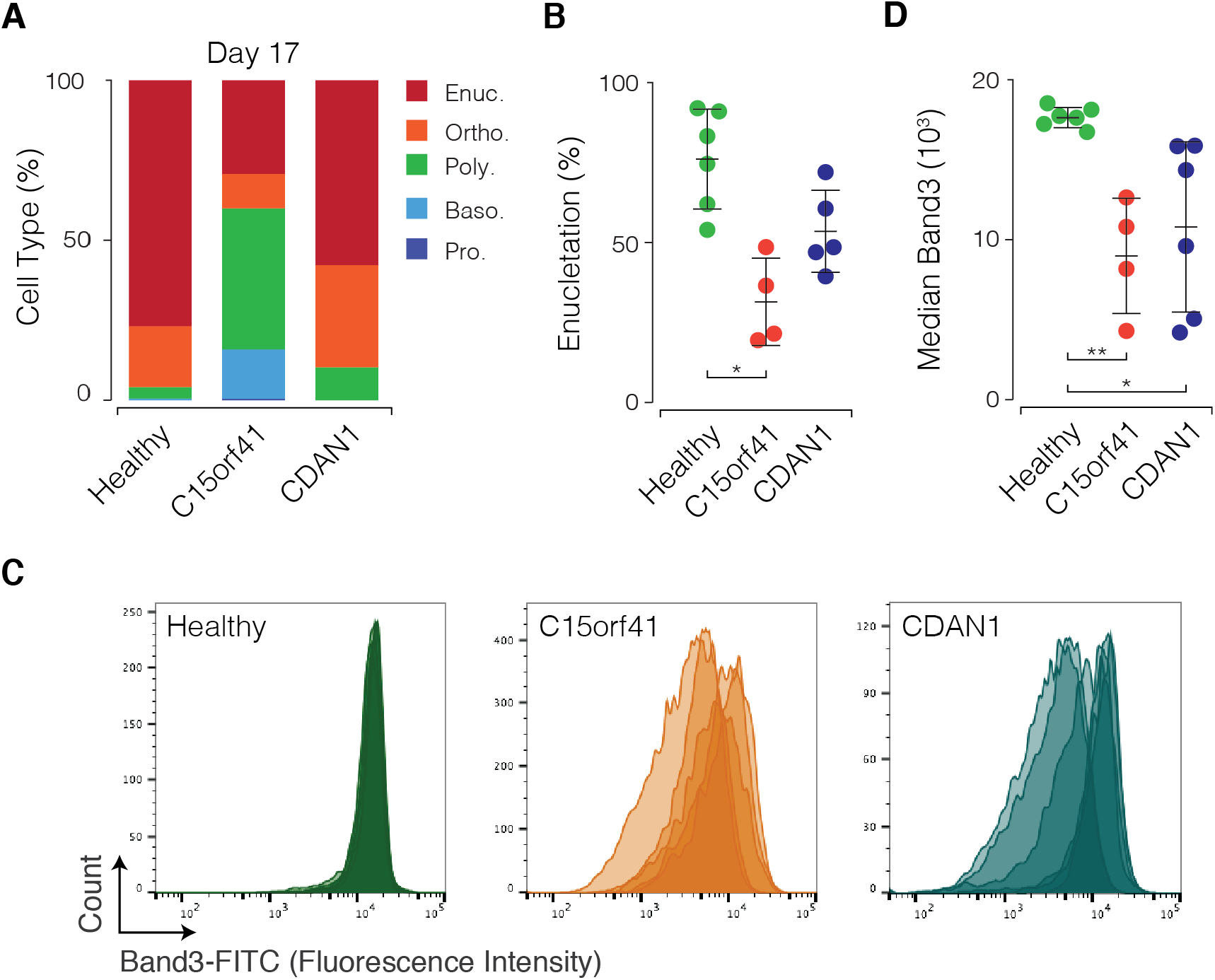
CDA-I erythroblasts show a delay in differentiation and reduced enucleation, most noticeable in *C15orf41* patients. (A) Percentages of erythroblast stages scored from stained cytospins at day 17 of *ex vivo* culture. (B) Percentage enucleation ±SD assessed by counts from cytospins at day 17 of cultured erythroblasts from healthy donors (n=6), patients with mutations in *C15orf41* (n=3) and patients with mutations in *CDAN1* (n=4). (\**P*=.049 with Kruskal-Wallis test) (C) FACS histograms of Band 3 intensity (in the CD235a^+^ population) at day 17 (healthy donors n=6, *C15orf41* patients n=3 and *CDAN1* patients n=4). (D) Median fluorescence intensity (MFI) ±SD of Band 3-FITC at day 17 (** *P*=.0088 and * *P*=.0127 with Kruskal-Wallis test

### Cultured erythroblasts can be used to diagnose CDA-I

The pathognomic diagnostic finding for CDA-I until now has been identification of specific heterochromatin abnormalities in bone marrow biopsies by TEM. The *ex vivo* culture system described here was used to confirm the diagnosis of CDA-I in a patient with a mutation in *CDAN1* (UPID16) without the need for an invasive bone marrow aspirate (Figure 4C). Instead, genetic analysis was conducted on the patient, who presented with unexplained anaemia, using the Oxford Red Cell Panel (ORCP)^34^ to identify any potentially pathogenic mutation (Figure 6). Following identification of a *CDAN1* variant, CD34^+^ progenitors from UPID16 were extracted and grown in our differentiation system. After 11 days in culture TEM was conducted on the resulting intermediate erythroblasts and revealed 29% of erythroblasts with abnormal chromatin morphology, thus confirming diagnosis of CDA-I.

**Figure 6:**
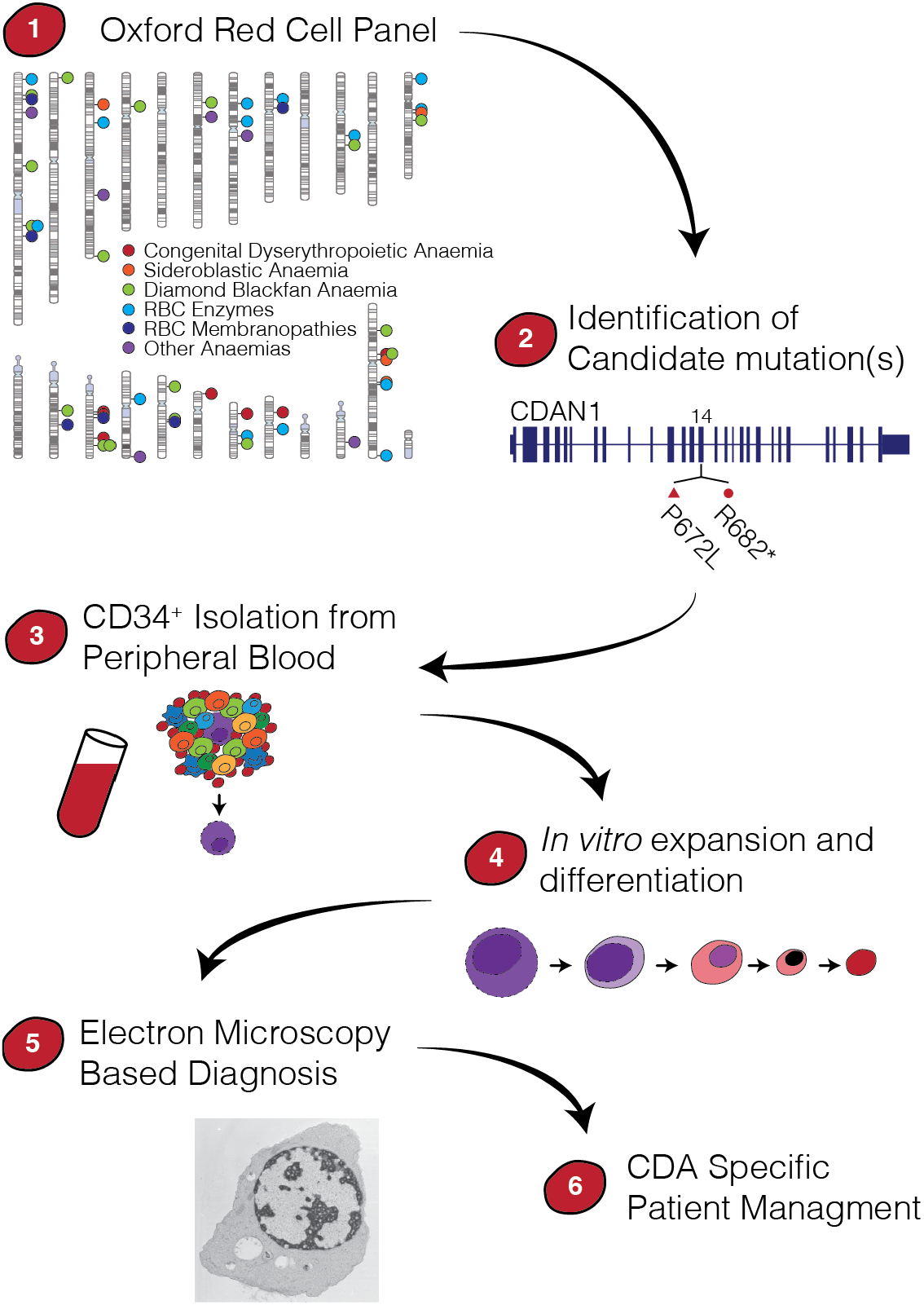
Overview of the strategy for patient (UPID16) diagnosis using cultured erythroblasts. Mutations in the two known causative genes were identified from gDNA of patient peripheral blood using a targeted re-sequencing Oxford Red Cell Panel^34^ and validated by confirmation of the presence of spongy heterochromatin, thus confirming the diagnosis of CDA-I.

## Discussion

Erythropoiesis is a complex, multi-step process coupling cell division with differentiation, starting from a heterogeneous population of haematopoietic stem cells^35^ and resulting in mature reticulocytes.^36^ Here we demonstrate that this can be modeled in an *ex vivo* culture system with erythroblasts passing through the expected stages of differentiation with appropriate expression of erythroid cell surface markers.^9^ By day 17 in our system approximately 70-80% of the cells were enucleated (Supplemental Figure 6B) indicating that erythroid differentiation was able to reach completion. Further, we were able to recapitulate the cardinal clinical feature of CDA-I, where we show by TEM that up to 40% of cultured CDA-I erythroblasts have spongy heterochromatin. In agreement with reported clinical findings^32,37^ we also see high levels of the hepcidin suppression factor GDF15 in cultured erythroblasts from CDA-I patients, again indicating that our culture system recapitulates common features of CDA-I and can therefore be used to elucidate mechanisms underlying the ineffective erythropoiesis in this disease.

While *ex vivo* culture from HSCs reproduces aspects of normal erythropoiesis and provides a source of erythroid cells to study, a number of confounding factors have been identified that make interpretation of the data difficult.^14,38^ Therefore, in order to investigate differences between patient-derived erythroblasts and those from healthy controls we took extreme care to stage the cultures using an array of methods that included FACS and CyTOF, which both rely on immunophenotyping cell populations. While such methods showed healthy and diseased erythroblasts were immunologically similar during the expansion phase and into early differentiation, aspects of disordered erythropoiesis became evident after this time by several parameters. Chromatin accessibility is a preferred approach for cell type classification over other methods such as RNA-seq and has been used to characterise all the blood lineages from bone marrow in humans.^31^ Further, it is well documented that subsets of regulatory elements are systematically activated and repressed to enable commitment to the different haematopoietic lineages and particularly during erythroid-megakaryocyte differentiation.^39^ Therefore we chose to use ATAC–seq as a genome-wide method of staging the cells. When our samples were aligned with a defined ATAC-seq erythroid trajectory,^31^ normal and patient material became clearly distinguishable indicating that CDA-I patients have a more precursor-like accessibility profile, with a disruption in progression through erythropoiesis leading to the reduced erythroid output we observe in CDA-I. In agreement, cytospins contained increased numbers of erythroid precursors and fewer enucleated cells. Although dysregulated Band 3 expression has been observed in MDS patients,^9^ the abnormal expression of Band 3 that we report has not been previously described in CDA-I patients and aligns with our other observations of affected terminal differentiation.

In terminal erythropoiesis, erythroid progenitors have a rapid cell cycle with a close association between the number of cell divisions and differentiation in order to generate the correct numbers of terminal erythroblasts.^40^ Proerythroblasts from umbilical cord CD34^+^ cells have been shown to have limited proliferative capacity through the successive stages to orthochromatic erythroblasts.^9^ Importantly, defects in various cell cycle genes have been reported to result in anaemia.^41^ Our data show, for the first time, increased proliferation of patient erythroblasts compared to healthy cells, which occurs in parallel with morphological and DNA accessibility differences that could impact on the closely coupled cell cycle progression and differentiation speed. Interestingly, expansion of the numbers of erythroid cell precursors, particularly the proerythroblasts, has been reported in ineffective erythropoiesis observed in patients with β-thalassemia.^30^ Further it has been shown that the tight link between the number of cell divisions and differentiation can be uncoupled by treatment with glucocorticoids, thought to be acting on developmental regulatory proteins.^42^ How this occurs in the case of CDA-I is not yet clear although previous findings from non-erythroid cell lines show Codanin-1 to be cell cycle regulated^7^ and cause an increase in cell cycle rate if over-expressed.^5^

The final stages of erythropoiesis involve chromatin and nuclear condensation prior to expulsion of the pyknotic nuclei by enucleation^36,43^ and this process is highly organized^44^ with chromatin condensation playing an important role.^45^ The abnormal spongy heterochromatin observed in CDA-I could have a significant impact on the usual processes that precede enucleation, such as the selective loss of histones. Remarkably, a substantial number of erythroblasts progress to enucleation without developing the catastrophic changes in chromatin compaction and organisation apparent in spongy nuclei. This implies that the effects of the aberrant proteins must reach a threshold within individual cells to produce the pathognomic phenotype.

Recently it has been reported that two distinct types of CDA-I exist (CDA-Ia MIM 224120 and CDA-Ib MIM 615631).^33^ The differential diagnosis is based on the levels of Hb and MCV, with the CDA-Ib patients (caused by *C15orf41* mutations) thought to be more severely affected, although the haematological data of our patient cohort (Figure 2B) shows there is overlap between the blood indices in CDA-I patients (excluding those regularly transfused or venesected) irrespective of the mutation. However, using our culture system, we observe an enucleation defect and significantly reduced levels of Band 3 expression in late erythroblasts from *C15orf41* patients, as compared to those with mutations in *CDAN1*. This suggests that there may indeed be a distinction based on patient genotype and further that the CDA-I phenotype may be more severe when arising from *C15orf41* mutations.

Precise diagnosis of rare congenital anaemias such as CDA-I may take a number of years since some dyserythropoietic features are often present in other types of anaemia. Patients suffering from CDA-I tend to present in childhood and adolescence and until recently, diagnosis has required assessment of bone marrow by electron microscopy. Previously our laboratory has shown the benefits of using a targeted next generation sequencing panel (NGS) together with morphological analysis to confirm and improve the rate and accuracy of diagnosis of CDA-I.^34^ However the pathogenicity of novel *C15orf41* or *CDAN1* variants identified cannot be determined without further evidence. We propose that the *ex vivo* culture system reported here can be used for such validation, as we have clearly shown the pathognomic feature of spongy heterochromatin in our cultured erythroblasts which would enable a diagnosis of CDA-I to be made from peripheral blood. Benefits include avoidance of invasive bone marrow aspirates and associated risks,^46^ as well as precise diagnosis for patients who have often had many years of clinical investigation.^34^ Ultimately this could streamline the diagnosis pathway and improve care for patients with this rare genetic form of anaemia, as well as providing patient-specific erythroblasts for research.

In this study we provide a detailed characterisation of CDA-I patient-specific erythroblasts, which have been modeled *ex vivo*. Results show that we can recapitulate aspects of the disease pathology seen in CDA-I, including high levels of cells with spongy heterochromatin as well as increased GDF15 expression. We also report here for the first time that CDA-I patient erythroblasts have elevated levels of proliferation, together with persistence of erythroid precursors. The abnormalities identified during late erythropoiesis in Band 3 expression and levels of enucleation most markedly affect patients with mutations in *C15orf41*, providing firm evidence for a genotype/phenotype correlation in CDA-I.

## Acknowledgements

We would like to thank all the CDA-I patients for providing blood samples. We would like to acknowledge the flow cytometry facility at the WIMM for providing cell analysis services and technical expertise. The facility is supported by the MRC HIU; MRC MHU (MC_UU_12009); NIHR Oxford BRC; Kay Kendall Leukaemia Fund (KKL1057), John Fell Fund (131/030 and 101/517), the EPA fund (CF182 and CF170) and by the WIMM Strategic Alliance awards G0902418 and MC_UU_12025. We would like to acknowledge the Electron Microscopy Facility at the Sir William Dunn School of Pathology for conducting the majority of the TEM and Raman Dhaliwal for help with the imaging.

Further support came from grants to the Wolfson Imaging Centre Oxford (Wolfson Foundation 18272, joint MRC/BBSRC/ EPSRC MR/K015777X/1, Wellcome Trust Multi-User Equipment 104924/Z/14/Z). We would like to acknowledge Giorgio Napolitani and Michalina Mazurczyk for their help in the mass cytometry facility at the WIMM for providing technical expertise and cell analysis services. The facility is supported by the MRC HIU core funded project, reference MC_UU_00008 and the Oxford Single Cell Biology Consortium (OSCBC). This work was supported by the charity Congenital Anaemia Network (CAN), the Medical Research Council MC_UU_12009/1 and a Wellcome Trust Strategic Award (106130/Z/14/Z).

## Authorship

Contributions: C.S & D.J.D. extracted CD34 cells from CDA-I patient blood and normal donors respectively. C.S., D.J.D, M.G. performed the experiments with help from A.A.O. R.S. and D.J.D. undertook the ATAC analysis. D.J.P.F and E.J. performed the electron microscopy. J.M.B. conducted the immunofluorescence. M.B. conducted the CyTOF and the data was analysed by R.B. Q.A.H, S.O, R.R, K.R. and N.R. were the clinicians responsible for the care of the of the CDA-I patients. C.S., D.J.D., J.H., C.B., and V.J.B. conceived and designed experiments. D.R.H. provided conceptual advice and clinical oversight. D.J.D. created the figures, C.S. and V.J.B. wrote the paper and all authors reviewed and critically edited the manuscript.

