## Supplemental figures for "Modelling erythropoiesis in congenital dyserythropoietic anaemia type I (CDA-I)"

**Supplementary Data**

Caroline Scott[1], Damien J. Downes[1], Jill M. Brown[1], Christian Babbs[1], Aude-Anais Olijnik[1], Matthew Gosden[1], Robert Beagrie[1], Ron Schwessinger[1], Christopher A. Fisher[1], Anna Rose[1], David J.P Ferguson[2], Errin Johnson[3], Quentin. A Hill[4], Steven Okoli[5], Raffaele Renella[6], Kate Ryan[7], Marjorie Brand[8], Jim Hughes[1], Noemi Roy[9], Douglas R. Higgs[1] and Veronica Buckle[1].

[1]Weatherall Institute of Molecular Medicine, John Radcliffe Hospital, Oxford University, Oxford, United Kingdom; [2]Ultrastructural Morphology Group, NDCLS, John Radcliffe Hospital, Oxford, United Kingdom; [3]Sir William Dunn School of Pathology, South Parks Road, Oxford, United Kingdom; [4]Leeds Teaching Hospital NHS Trust, United Kingdom; [5]Imperial College, The Commonwealth Building, The Hammersmith Hospital, Du Cane Road, London, United Kingdom; [6]Pediatric Hematology-Oncology Research Laboratory, CHUV-UNIL Lausanne Switzerland; [7]Department of Haematology, Manchester Royal Infirmary, Oxford Road, Manchester, United Kingdom; [8]Sprott Center for Stem Cell Research, Ottawa Hospital Research Institute, Ottawa, Canada and [9]Department of Haematology, Oxford University Hospitals NHS Trust, Churchill Hospital, Old Road, Headington, Oxford, United Kingdom.

Supplemental Figure 1

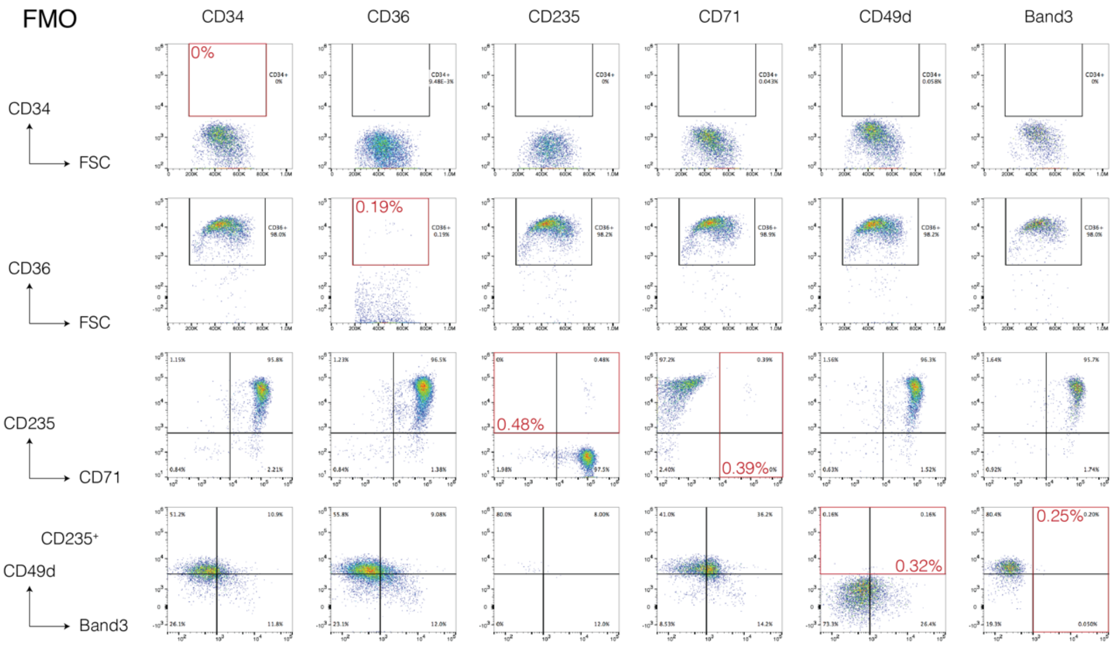


**Supplemental Fig 1. Fluorescence minus one (FMO) controls for FACS analysis.** Gating strategy for FACS analysis. Gates were set for each population on FMO. Red boxes indicated postive population for each marker with the percent of cells in this gate from the FMO shown.

Supplemental Figure 2

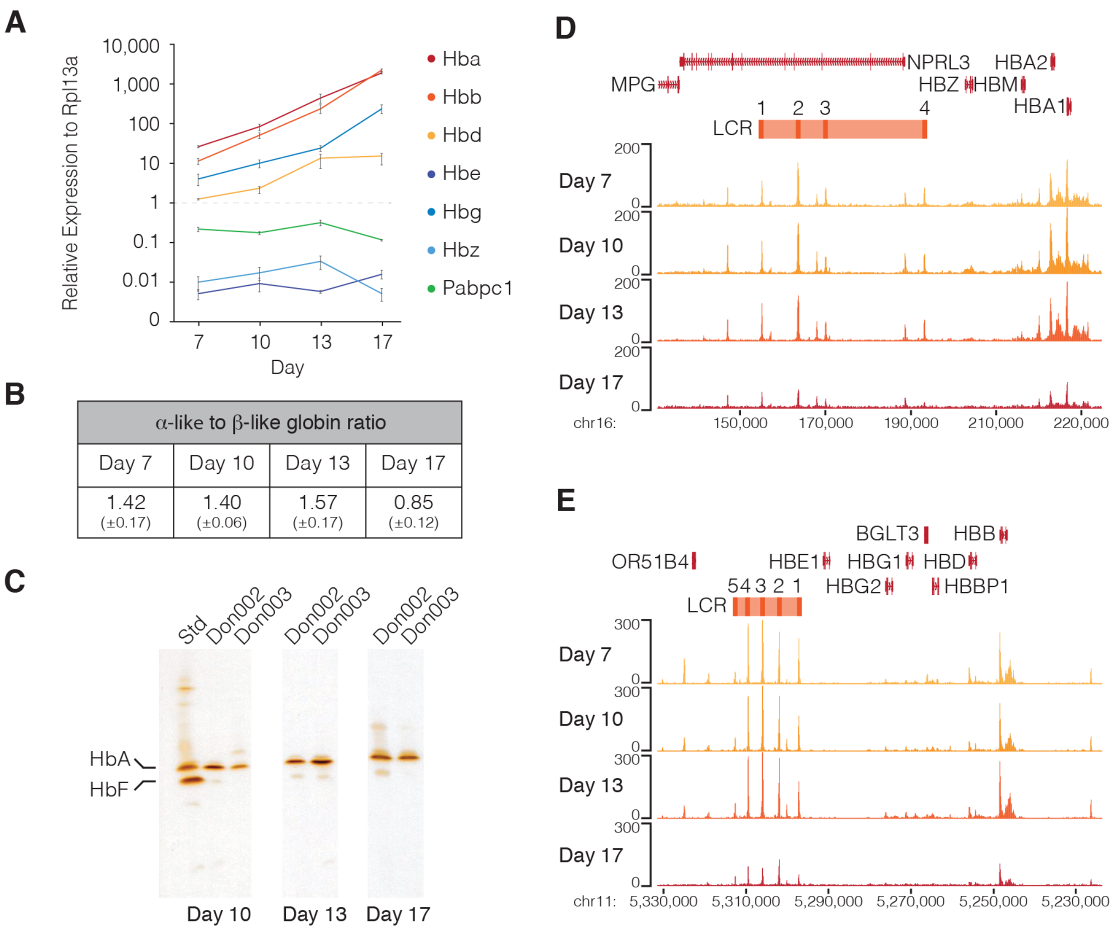


| α-like to β-like globin ratio | | | |
|---|---|---|---|
| Day 7 | Day 10 | Day 13 | Day 17 |
| 1.42 (±0.17) | 1.40 (±0.06) | 1.57 (±0.17) | 0.85 (±0.12) |

**Supplemental Fig 2. Cultured erythroblasts show an increase in DNA accessibility and expression of the adult globin genes during differentiation.** (A) Gene expression analysis of globins (mean ± SEM), throughout the differentiation normalized to RPL13A. Pabpc1 was used as a housekeeping gene. (B) Ratios of the alpha-like to beta-like globins (mean ± SEM) during erythroid differentiation. (C) IEF of erythroblasts from 2 healthy donors (Don002 and 003) at three timepoints during *ex vivo* differentiation. HbA is adult haemoglobin and HbF is foetal haemoglobin. (D) ATAC-seq of the alpha-globin locus at four time-points throughout *ex vivo* differentiation. (E) ATAC-seq of the beta-globin locus at four time-points throughout *ex vivo* differentiation.

Supplemental Figure 3

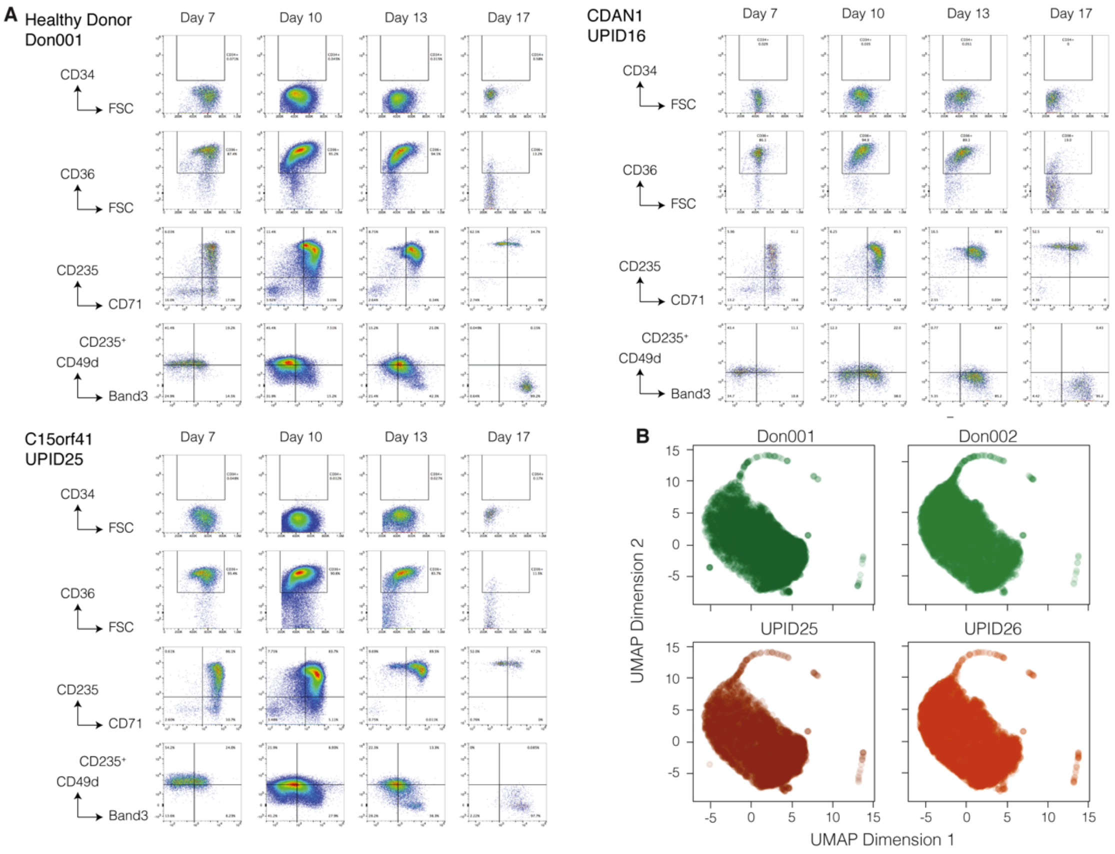


**Supplemental Fig 3. CDA-I patients and healthy donors show similar progression through erythropoiesis.** (A) Representative FACS profiles of cultured erythroblasts from a healthy donor, CDA-I patient with a mutation in *C15orf41* (UPID25) and CDA-I patient with a mutation in *CDAN1* (UPID16) during *ex vivo* differentiation. (B) UMAP plots of CyTOF data showing expression of 25 transcription factors and erythroid cell surface markers at day 11 of differentiation in healthy controls (Don001 & Don002) and CDA-I patients with mutations in *C15orf41* (UPID 25 & 26).

Supplemental Figure 4

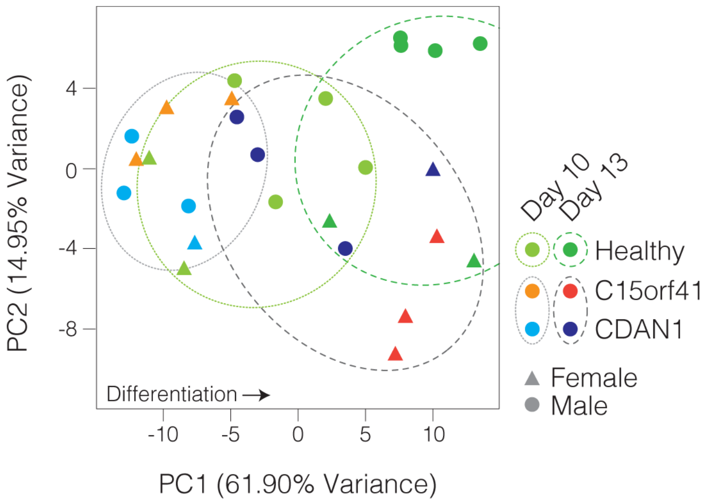


**Supplemental Fig 4. ATAC-seq analysis reveals a slight lag in differentiation of patient-derived erythroblasts.** Principle component analysis (PCA) of ATAC-seq. The distribution of cells along PC1 follows differentiation stage.

Supplemental Figure 5

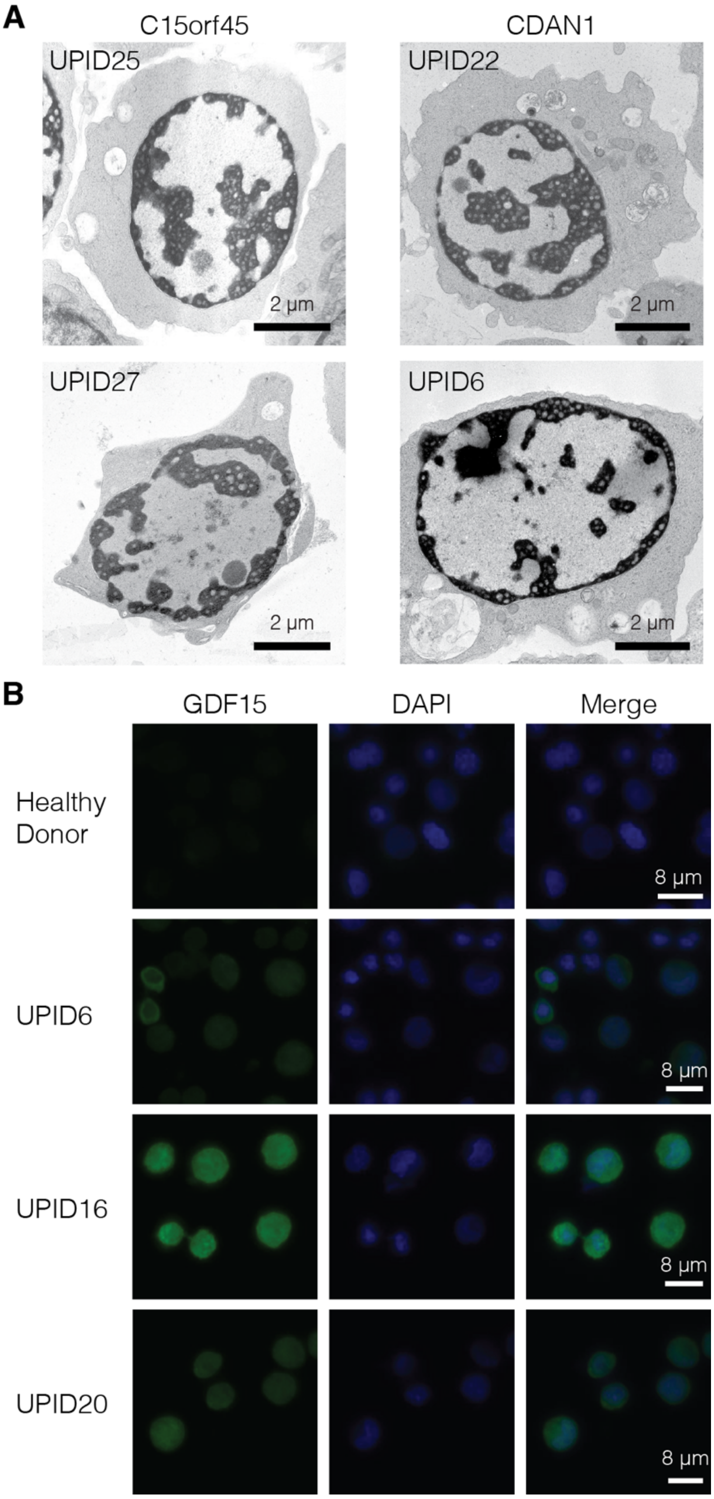


**Supplemental Fig 5. Ex vivo cultured patient erythroblasts show the diagnostic features of CDA-I.** (A) Representative examples of electron micrographs of abnormal nuclei seen in patients with mutations in *C15orf41* (n=2) and *CDAN1* (n=2). (B) Immunofluorescence of day 10 cultured erythroblasts from a healthy donor and *CDAN1* patients (UPID6, 16 and 20). GDF15 is detected with Alexa488 and

DAPI was used as a nuclear counter stain. The merged images are shown in the right hand panel.

Supplemental Fig 6.

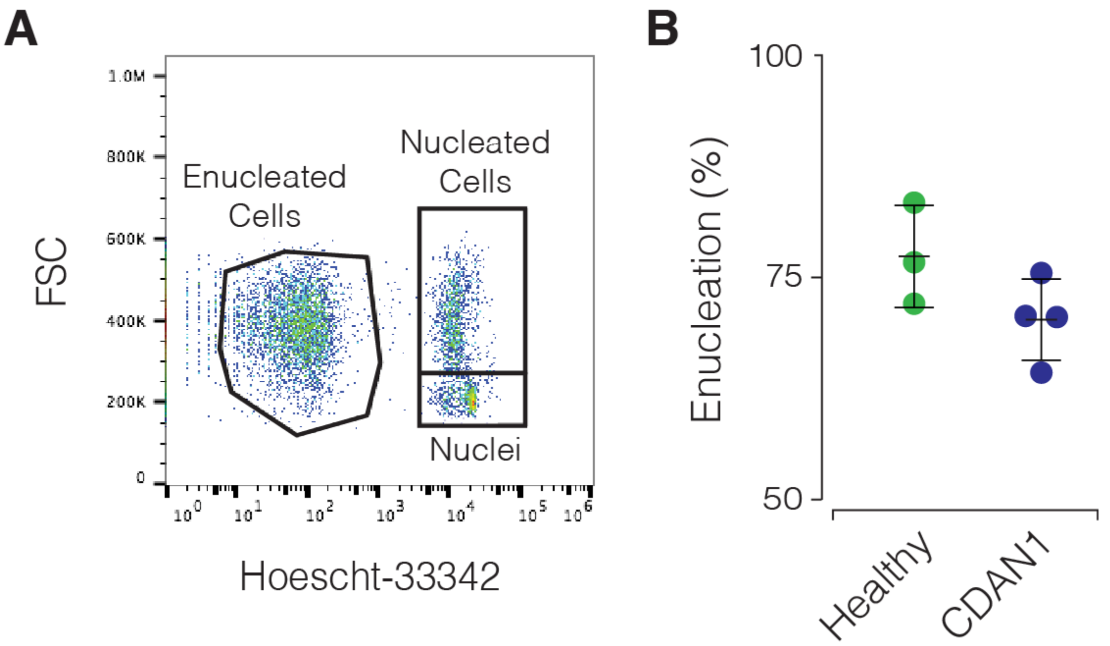


**Supplemental Fig 6. Levels of enucleation by FACS.** (A) Representative example of FACS analysis to determine enucleation at day 17 of *ex vivo* culture. (B) Percentage enucleation ± SD assessed by FACS in day 17 cultured erythroblasts from healthy donors (n=3) and patients with mutations in *CDAN1* (n=4).

**Summary of Supplemental Tables**

**Supplemental Table 1. Fluorophore conjugated antibodies used for staging the erythroid differentiation by FACS.**

**Supplemental Table 2: Taqman probes used for globin expression analysis.**

**Supplemental Table 3: Next generation sequencing depth.**

**Supplemental Table 4: Bed file of top 1000 ATAC peaks used for PCA plots.**

**Supplemental Table 5: Bed file of ATAC-seq non-TSS nucleosome depleted regions (NDR) used for PCA plots.**

**Supplemental Table 6: Pre-conjugated antibodies used for CyTOF.**
